# Transcriptome-scale analysis uncovers conserved residues in the hydrophobic core of the bacterial RNA chaperone Hfq required for small regulatory RNA stability

**DOI:** 10.1101/2024.09.05.607192

**Authors:** Josh McQuail, Miroslav Krepl, Kai Katsuya-Gaviria, Aline Tabib-Salazar, Lynn Burchell, Thorsten Bischler, Tom Gräfenhan, Paul Brear, Jiří Šponer, Ben F. Luisi, Sivaramesh Wigneshweraraj

## Abstract

The RNA-chaperone Hfq plays crucial roles in bacterial gene expression and is a major facilitator of small regulatory RNA (sRNA) action. The toroidal molecular architecture of the Hfq hexamer contains three well characterised surfaces which allow it to bind sRNAs to stabilise them and engage target transcripts. Hfq-interacting sRNAs are categorised into two classes based on the surfaces they use to bind Hfq. By characterising a systematic alanine mutant library of Hfq to identify amino acid residues that impact survival of *Escherichia coli* experiencing nitrogen starvation, we corroborated the important role of the three RNA binding surfaces for Hfq function. Surprisingly, we uncovered two conserved residues, V22 and G34, in the hydrophobic core of Hfq, to have a profound impact on Hfq’s RNA binding activity *in vivo*. Transcriptome-scale analysis revealed that V22A and G34A Hfq mutants cause widespread destabilisation of both sRNA classes. However, the alanine substitutions at these residues had no measurable impact on protein stability, structure or equilibrium binding to target sRNAs *in vitro*. We propose that V22 and G34 are key to the cooperative function among the RNA-binding surfaces of Hfq, a mechanism especially critical under cellular conditions when there is an increased demand for Hfq.

## Introduction

The post-transcriptional regulation of RNA stability and translational efficiency allows enormous flexibility in the control of genetic information in bacteria. Small non-coding regulatory RNA molecules, called sRNAs, play a pivotal role in post-transcriptional regulation of messenger RNA (mRNA). sRNAs determine whether the targeted mRNAs are destined for translation, translational repression, or degradation. A core component of post-transcriptional regulation are specialised RNA binding proteins which facilitate the interaction between sRNAs and their cognate target mRNAs.

In many bacteria, the majority of the interaction between sRNA and mRNA involve the RNA binding protein Hfq (1,2). However, other functions of Hfq in regulating translation and degradation of mRNAs independently of the sRNA-mediated regulatory pathway have also been documented (3–7). Hfq is a member of the Sm/LSm superfamily of RNA-binding proteins, the other members of which can be found in almost every organism from all three domains of life (8). These proteins have a common structural ‘core’ comprised of an N- terminal α-helix followed by a twisted five-stranded β-sheet and assemble into a variety of quaternary structures ranging from pentamers to octamers (9). Structures of the Hfq conserved ‘core’ region from several different bacterial species are available, and in all cases a hexameric ring structure has been observed (10–13). The conserved core of the Hfq monomer consists of an α–β_1–5_ fold, and when these pack into a hexamer, three faces are presented for interaction with RNA. The ‘proximal face’ is close to the amino-terminal of Hfq, and the ‘distal face’ lies on the opposite side of the Hfq hexamer, close to the carboxyl-terminal of Hfq; the ‘rim region’, which separates the proximal and distal faces provides additional RNA binding sites. Appended to the conserved core is a structurally disordered carboxyl-terminal domain (CTD) that is variable in size and sequence. The CTD of Hfq in *E. coli* acts synergistically with the other RNA binding faces on the conserved core and contributes to the specificity of its RNA annealing activity (14–16).

The biogenesis of sRNAs can occur in multiple ways. sRNA can be made by transcription of a stand-alone noncoding gene; or can be derived from coding genes by premature transcription termination in 5′ untranslated region (UTR), by transcription starting from inside the coding region of a gene but using the same terminator (31-UTR derived) or by processing of mRNA by RNases. In all cases, interaction with Hfq is important for the intrinsic stability of sRNAs and mutations in RNA interacting surfaces on Hfq can lead to decreased sRNA stability (17). Typically, amino acid (aa) substitutions at the conserved surface exposed proximal residue Q8, rim residues R16 and R17 and proximal face residues Y25 and K31 are widely used in many studies to understand Hfq interaction with RNA. An elegant study by Schu et al (17) utilised these Hfq mutants with 24 different sRNAs and their cognate mRNA partners to describe two different interaction modes of sRNA to Hfq (designated as class I and class II), which involve cooperation of the different RNA binding surfaces on Hfq. Class I sRNAs interact with the proximal face and rim of the Hfq ring, whereas their mRNA targets bind the distal face of the ring; class II sRNAs bind to the proximal and distal faces and base pair with mRNA targets that interact with the rim (Figure 1A) (17). Further, Malecka et al revealed that rim region and the distal face are important for compacting mRNA for optimal binding to cognate Hfq-bound sRNA (18), further underscoring that the cooperation between the RNA binding surfaces is crucial for Hfq function.

**Figure 1.**
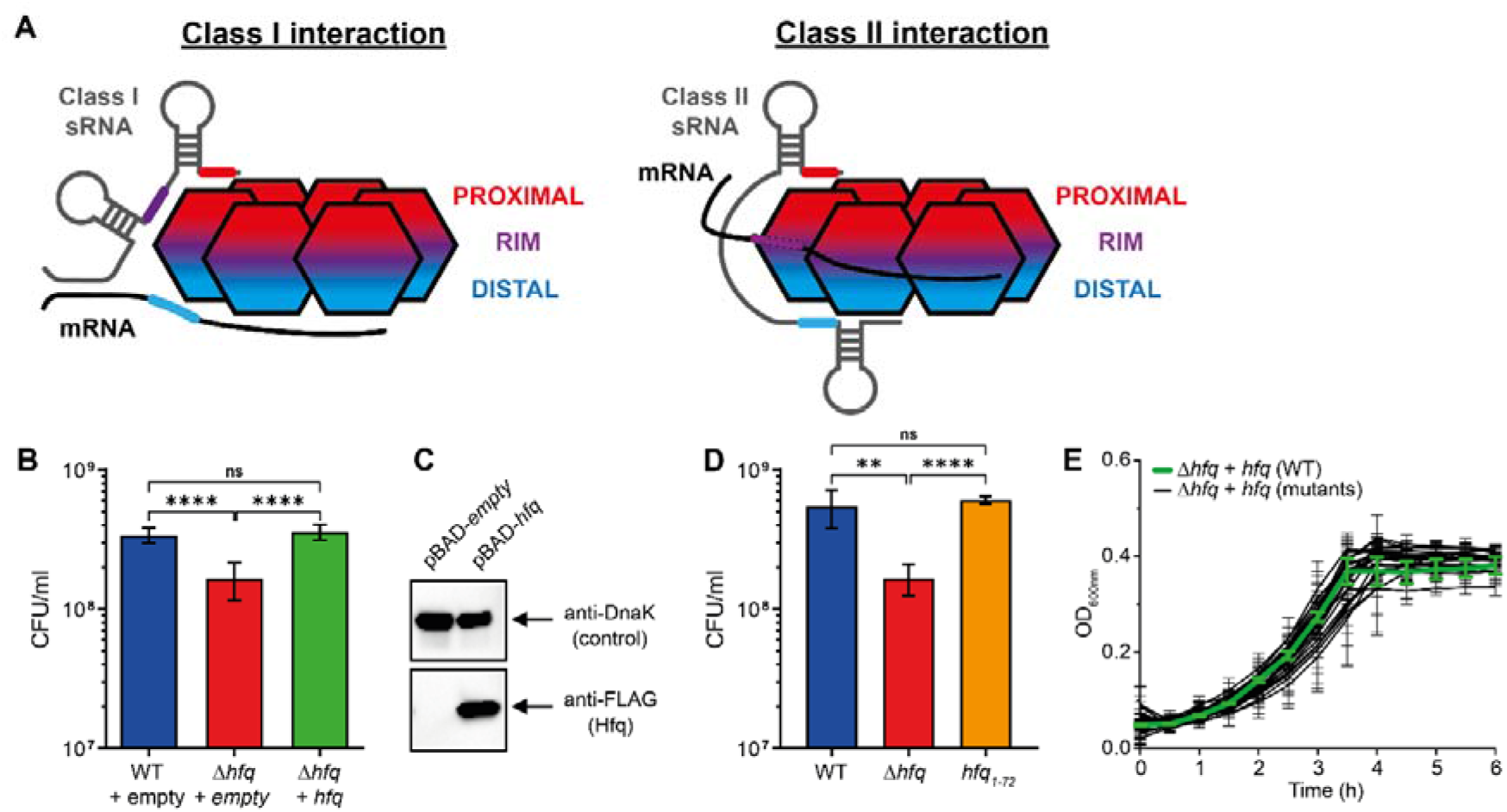
Hfq is required for survival during N starvation in *E. coli* **(A)** Schematic showing the established mechanisms of interaction between class I and class II sRNA with Hfq and their target mRNA. Hfq monomers are represented by the six hexagons, the red section represents the proximal RNA-binding face, the purple section represents the rim region, and the blue section represents the distal RNA-binding face. Class I and class II sRNA are represented by the grey lines, and mRNA are represented by black lines. Regions of either sRNA or mRNA which associate with the different RNA-binding surfaces of Hfq are coloured accordingly. **(B)** Viability of WT and Δ*hfq E. coli* expressing plasmid-borne Hfq (pBAD24-*hfq*-3xFLAG) measured by counting CFU at N-24. **(C)** Representative immunoblot of whole-cell extracts of Δ*hfq E. coli* containing either a pBAD18 empty vector control, or pBAD24-*hfq*-3xFLAG, sampled at N-24. The immunoblots were probed with anti-FLAG antibody and anti-DnaK antibody (loading control). **(D)** Viability of WT and Δ*hfq E. coli,* and *E. coli* expressing Hfq with a C-terminal truncation from residues 73-102 (*hfq_1-72_*) measured by counting CFU at N-24. **(E)** Growth of Δ*hfq E. coli* expressing plasmid-borne WT or alanine mutants of Hfq (pBAD24-*hfq*-3xFLAG) under N limiting conditions. In **(B)**, **(D)** & **(E)** error bars represent standard deviation (n = 3). In **(B)**, & **(D)** statistical analysis performed by Brown-Forsyth and Welch’s ANOVA. (****, P<0.0001; **, P<0.01).

Recently, we showed that in *E. coli* experiencing N starvation, the Hfq-mediated post-transcriptional regulation is extensive, dynamic and important for cell survival (19,20). Despite the central role Hfq has in sRNA stabilisation and post-transcription regulation of gene expression, the involvement of individual aas of Hfq in its function has never been systematically investigated. In this study, we characterised a systematic alanine mutant library of Hfq under N starvation and provide new transcriptome-wide insights into sRNA binding and stabilisation by Hfq under *in vivo* conditions when there is an increased demand for Hfq.

## Materials and methods

### Construction of Hfq mutants and bacterial growth conditions

A systematic alanine mutant library of Hfq was created by site-directed PCR mutagenesis using pBAD24-*hfq*-3xFLAG (20) as the template and appropriate primers (Supplementary Data 1) and confirmed by DNA sequencing. Note that, due to the cloning procedure, an additional V aa residue was introduced at aa position 2 of Hfq so that the N-terminal aa sequence of Hfq in pBAD24-*hfq*-3xFLAG is MVAK instead of MAK. The *E. coli* strain BW25113 was used for all the experiments. The WT and Δ*hfq* BW25113 strains were obtained from the *E. coli* Genetic Stock Center. The construction of the *hfq*_1-72_ BW25113 strain is described in (20). N starvation experiments were conducted as described in (21). Briefly, unless otherwise stated bacteria were grown in 10 mM NH_4_Cl (for overnight cultures) or 3 mM NH_4_Cl (for N starvation experiments) in Gutnick minimal medium (33.8 mM KH_2_PO_4_, 77.5 mM K_2_HPO_4_, 5.74 mM K_2_SO_4_, 0.41 mM MgSO_4_), supplemented with Ho-LE trace elements (22), 0.4% (w/v) glucose and 100 μg/ml of ampicillin at 37°C in a shaking (700 rpm) SPECTROstar OMEGA plate reader (BMG LABTECH)(21). To induce expression of Hfq, L-arabinose was added at final concentration of 0.2% (w/v). Growth of bacterial cultures was measured either by optical density at 600 nm (OD_600_) over time or determined by measuring colony forming units (CFU) from serial dilutions on lysogeny broth (LB) agar plates.

### Immunoblotting

Whole cell extracts of N-24 bacteria were prepared by pelleting bacterial cell from a 1 ml culture and resuspension in denaturing-PAGE loading buffer. Following denaturing-PAGE of the sample, the gel was transferred onto polyvinylidene difluoride (PVDF) membrane (0.2 μm). For immunoblotting of Hfq protein, mouse monoclonal Anti-FLAG^®^ M2 antibody (Merck, F1804) was used at 1:1000 dilution. For the loading control, mouse monoclonal anti-DnaK antibody (Enzo, 8E2/2) was used at 1:1000 dilution. HRP sheep anti-mouse IgG (GE healthcare, NA931) at 1C:C10,C000 dilution as the secondary antibody. ECL Prime Western blotting detection reagent (GE Healthcare, RPN2232) was used to develop the blots, which were analysed on the ChemiDoc MP imaging system and bands quantified using Image Lab software.

### Preparation of the molecular dynamics (MD) calculations

Chain A of the *E. coli* Hfq hexamer X-ray structure (PDB: 1HK9) (23) was used to obtain the initial coordinates of the Hfq monomer in all simulations. To increase sampling efficiency, the flexible C-terminal residues (aas 66-69) were removed from the X-ray structure of Hfq (23). The F11A, L12A, V22A, G34A, I24A, I36A, and Y55A mutants were obtained by substituting the individual residues in the WT structure. The topology and coordinate files for MD simulations were created with the xLeap program of the AMBER 20 package (24). To describe the protein, we used either the ff19SB or ff12SB protein force fields. The ff12SB is the earlier version of the ff14SB (25) which was however shown to provide superior description for some systems (26). The ff19SB force field is the successor to both of these older force fields, showing improvements for proteins with disorder and large-scale dynamics (27). To describe the water molecules, we have used the OPC (28) and SPC/E (29) water models in the ff19SB and ff12SB simulations, respectively, as recommended for these protein force fields (25,27). In all simulations, the protein was surrounded in an octahedral box of water molecules with minimal distance of 12 Å (SPC/E) or 13 Å (OPC) between the solute and the box border. Physiological ion concentration of 0.15 M was established by adding KCl ions (30) at random positions around the solute. For each system, we have performed minimization and equilibration with pmemd.MPI module of AMBER 20 according to the standard protocol (31).

### Standard MD simulations

Standard (unbiased) MD simulations were run for 5 μs, using the pmemd.cuda module and RTX 2080ti GPUs. Selected trajectories were extended up to 20 μs and multiple trajectories of each system were obtained (Table 1). The SHAKE algorithm (32) along with the hydrogen mass repartitioning (33) were applied in all simulations, allowing a 4 fs integration step. The long-range electrostatics was treated using the particle mesh Ewald methodology (34) and periodic boundary conditions were imposed to handle the system border bias. The cut-off distance for Lennard-Jones interactions was 9 Å. Langevin thermostat and Monte Carlo barostat (24) were used to maintain the temperature and pressure around 300 K and 1 bar, respectively.

**Table 1.**
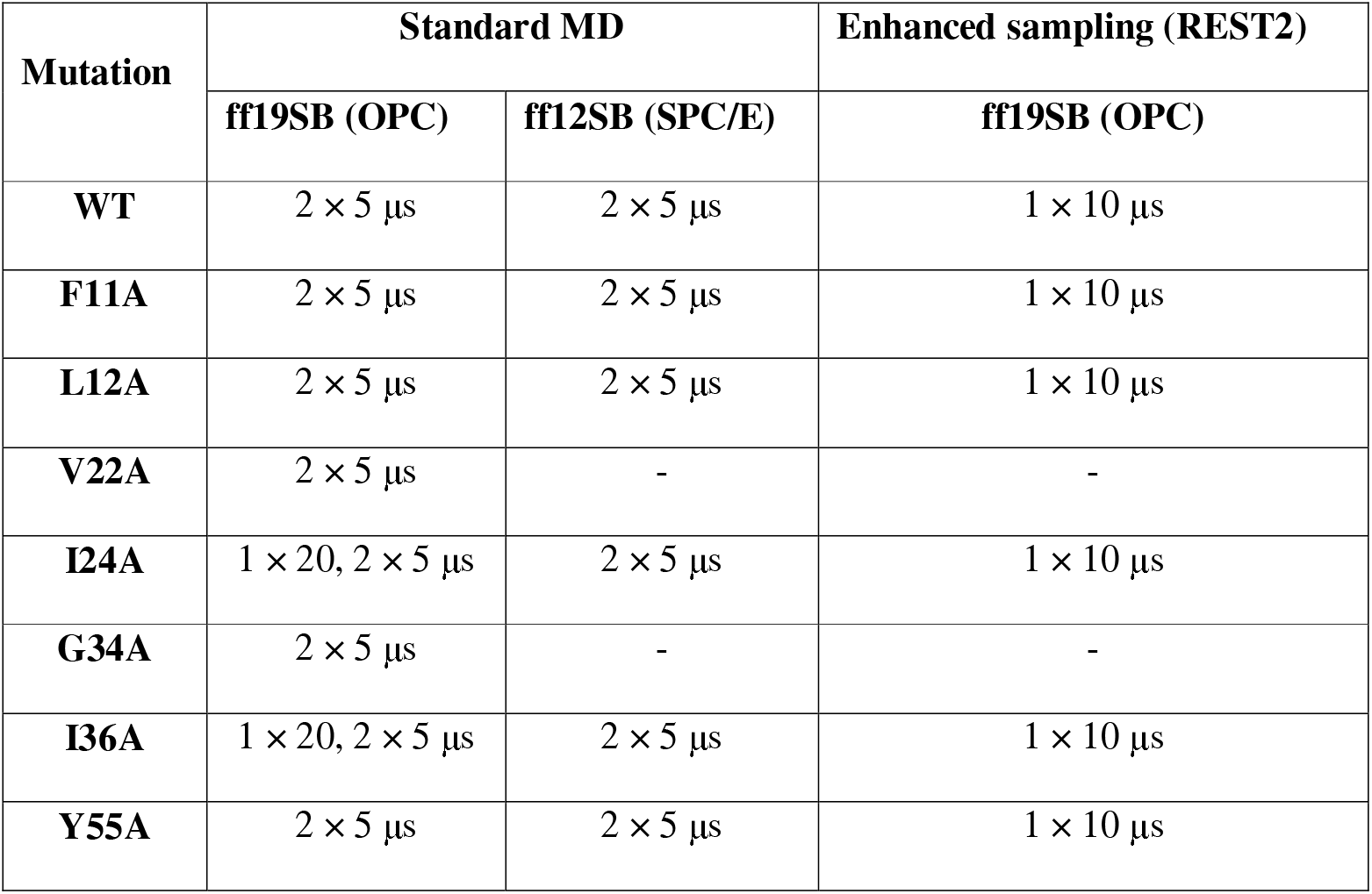
List of molecular dynamics simulations.

| **Mutation** | **Standard MD** |  | **Enhanced sampling (REST2)** |
| --- | --- | --- | --- |
|  | **ff19SB (OPC)** | **ff12SB (SPC/E)** | **ff19SB (OPC)** |
| **WT** | $2 \times 5 \mu\text{s}$ | $2 \times 5 \mu\text{s}$ | $1 \times 10 \mu\text{s}$ |
| **F11A** | $2 \times 5 \mu\text{s}$ | $2 \times 5 \mu\text{s}$ | $1 \times 10 \mu\text{s}$ |
| **L12A** | $2 \times 5 \mu\text{s}$ | $2 \times 5 \mu\text{s}$ | $1 \times 10 \mu\text{s}$ |
| **V22A** | $2 \times 5 \mu\text{s}$ | - | - |
| **I24A** | $1 \times 20, 2 \times 5 \mu\text{s}$ | $2 \times 5 \mu\text{s}$ | $1 \times 10 \mu\text{s}$ |
| **G34A** | $2 \times 5 \mu\text{s}$ | - | - |
| **I36A** | $1 \times 20, 2 \times 5 \mu\text{s}$ | $2 \times 5 \mu\text{s}$ | $1 \times 10 \mu\text{s}$ |
| **Y55A** | $2 \times 5 \mu\text{s}$ | $2 \times 5 \mu\text{s}$ | $1 \times 10 \mu\text{s}$ |

### Enhanced-sampling REST2 simulations

For each system, we performed REST2 (Replica Exchange with Solute Tempering) simulations (35) with 12 replicas, with the scaling factor (lambda) ranging from 1 to ∼0.6. The REST2 is a “brute-force” enhanced sampling method that does not follow any particular user-defined collective variable (CV). Instead, the accelerated sampling is achieved by modifying the potential energy function of the system, lowering the energy barriers for conformational transitions of the protein. This is coupled with the replica exchange protocol.

The REST2 can be used to observe conformational changes that are beyond the timescale of standard MD simulations. The main disadvantage of this protocol is loss of the information on kinetics. In all of our REST2 simulations, the entire Hfq monomer was scaled (i.e., included in the hot zone). Only the ff19SB force field was utilized for REST2 and the CMAP potentials were scaled by the same factor as the dihedral potentials. The lambda values across the replica ladder were chosen for each system to maintain average successful exchange rate of 25%. The REST2 simulations were performed using the pmemd.cuda.MPI module of AMBER 20 and their length was 10 μs (Table 1). The simulation settings were the same as for the standard simulations (see above) except the production phase of the REST2 simulations was performed in constant volume ensemble.

### Analyses of the simulations

We used cpptraj and VMD for analysis and visualization of all MD trajectories (36,37). Graphs and molecular figures were prepared with gnuplot and povray, respectively. The native contacts analysis was performed with cpptraj, by first compiling a list of interatomic distances shorter than 7 Å present in the initial structure, i.e., the native contacts. These distances were subsequently evaluated at every tenth frame of all trajectories, with the native contact counted as present for distances shorter than 7 Å. Only the distances between heavy atoms of residues at least four amino acids apart in the sequence were considered for the analysis. The native contacts analysis was performed on second halves of the standard MD trajectories of each system, with multiple trajectories combined into a single ensemble. Due to slightly different overall number of native contacts in each mutant, we present percentages of the native contacts preserved rather than their absolute numbers. For the REST2 simulations, the analysis was performed on the second half of the reference (unscaled) replica of each system. The resulting data was used to calculate histograms, with bin size set to 50 and the distributions normalized. All the trajectories were extensively visually inspected.

### Hfq purification

Hfq variants were expressed from pBAD vector in Δ*hfq* TOP10 cells grown at 37°C, induced with 0.1 % L-arabinose at OD = 0.4-0.6 and harvested by centrifugation after an overnight expression at 18°C. Cells were resuspended in Lysis Buffer (50 mM Tris-HCl pH 8.0, 1.5 M NaCl, 250 mM MgCl_2_, cOmplete™ EDTA-free Protease Inhibitor Cocktail [Roche]). Resuspended cells were lysed using Emulsiflex C5 (Avestin) high-pressure homogeniser (1000 bar). Lysate was clarified by centrifugation (4°C, 37500g, 30 mins). Supernatant was incubated at 85°C for 10 minutes and centrifuged (20°C, 37500g, 30 mins). Resulting supernatant was incubated with ammonium sulfate (0.9M) and centrifuged (20°C, 37500g, 30 mins). Supernatant was filtered (Sartorius Minisart® 0.45 μm) and loaded on 5 mL HiTrapButyl HP (Cytiva), previously equilibrated with Butyl Buffer A (50 mM Tris pH 8.0, 1.5 M NaCl, 1 M (NH_4_)_2_SO_4_). Proteins were eluted with an isocratic gradient of Butyl Buffer B (50 mM Tris pH 8.0). Fractions containing Hfq were pooled and diluted 1:5 with Butyl Buffer B. Pooled sample was then loaded on 5 mL HiTrap Heparin HP (Cytiva), previously equilibrated with Heparin Buffer A (50 mM Tris-HCl pH 8.0, 100 mM NaCl, 100 mM KCl), and eluted with an isocratic gradient of Heparin Buffer B (50 mM Tris-HCl pH 8.0, 1 M NaCl, 100 mM KCl). Fractions containing Hfq were pooled and concentrated (Amicon® Ultra-15 30 KDa, Millipore). Protein solution was loaded on Superdex 200 Increase 10/300 GL size exclusion column (S200; Cytiva) previously equilibrated with Hfq Storage Buffer (50 mM Tris-HCl pH 8.0, 100 mM NaCl,100 mM KCl, 5 % glycerol).

### Circular Dichroism (CD) spectroscopy

Measurements were made with an Aviv 410 CD spectrometer. Spectra were measured using 1 mm pathlength quartz cuvettes. Spectra in the near-UV range are averages of 10 measurements, and the far-UV are averages of 5 spectra. The average spectra were normalised for sample absorbance at 280 nm.

### Crystallisations, X-ray diffraction data collection and model refinement

Crystals were prepared of the pA_4_/Hfq V22A mutant using a mosquito crystallisation robot, and crystals were treated briefly with cryoprotectant, flash frozen and stored in liquid nitrogen. Diffraction data were collected at Diamond Light source station IO4 (mx33658-55) at wavelength 0.954 angstrom, and intensities were integrated and scaled in automated mode using xia2 (Table 2). The cell dimensions were indexed as P1 a=69.1819(12) b=69.2115(10) c=73.193(3) alpha 63.542(2) beta 89.304(2) gamma 60.0427(12), which is a pseudo-hexagonal setting. The structure was solved using molecular replacement the core residues 6- 66 of *E.coli* Hfq, based on PDB enbtry 3GIB. Two Hfq hexamers occupy the asymmetric unit, leaving 38% solvent contact, and the hexamers stack tightly through interactions of the proximal face and circumferential rim, but leave gaps between the stacks on the proximal face. The space in the lattice is likely filled with the N- and C-terminal tails, but these are comparatively disordered. The diffraction was anisotropic, which may reflect the disorder due to the packing gaps. One molecule of pA_4_ could be traced at the proximal face of one of the hexamers, and disordered portions of the RNA polymer could be fit at the proximal surface of the other hexamer. The model was refined using phenix refine (38), refmac (39) and extensive model building with coot (Table 3) (40). Anomalous fouriers did not reveal any peaks that may be due bound metal.

**Table 2.**
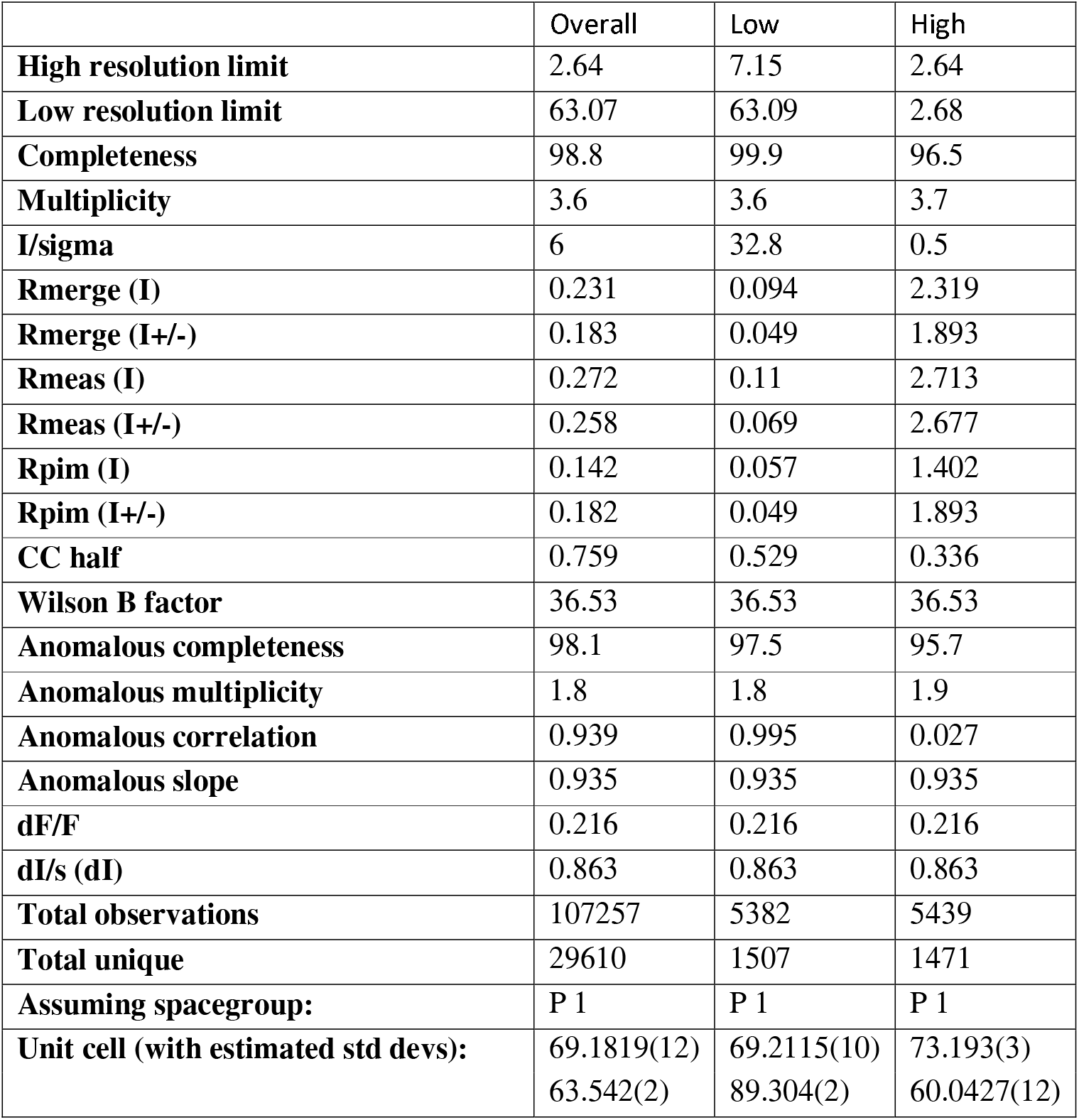
Crystallography Parameters.

|  | Overall | Low | High |
| --- | --- | --- | --- |
| **High resolution limit** | 2.64 | 7.15 | 2.64 |
| **Low resolution limit** | 63.07 | 63.09 | 2.68 |
| **Completeness** | 98.8 | 99.9 | 96.5 |
| **Multiplicity** | 3.6 | 3.6 | 3.7 |
| **I/sigma** | 6 | 32.8 | 0.5 |
| **Rmerge (I)** | 0.231 | 0.094 | 2.319 |
| **Rmerge (I+/-)** | 0.183 | 0.049 | 1.893 |
| **Rmeas (I)** | 0.272 | 0.11 | 2.713 |
| **Rmeas (I+/-)** | 0.258 | 0.069 | 2.677 |
| **Rpim (I)** | 0.142 | 0.057 | 1.402 |
| **Rpim (I+/-)** | 0.182 | 0.049 | 1.893 |
| **CC half** | 0.759 | 0.529 | 0.336 |
| **Wilson B factor** | 36.53 | 36.53 | 36.53 |
| **Anomalous completeness** | 98.1 | 97.5 | 95.7 |
| **Anomalous multiplicity** | 1.8 | 1.8 | 1.9 |
| **Anomalous correlation** | 0.939 | 0.995 | 0.027 |
| **Anomalous slope** | 0.935 | 0.935 | 0.935 |
| **dF/F** | 0.216 | 0.216 | 0.216 |
| **dI/s (dI)** | 0.863 | 0.863 | 0.863 |
| **Total observations** | 107257 | 5382 | 5439 |
| **Total unique** | 29610 | 1507 | 1471 |
| **Assuming spacegroup:** | P 1 | P 1 | P 1 |
| **Unit cell (with estimated std devs):** | 69.1819(12)<br>63.542(2) | 69.2115(10)<br>89.304(2) | 73.193(3)<br>60.0427(12) |

**Table 3.** Crystallography data collection and refinement statistics. (Statistics for the highest-resolution shell are shown in parentheses).

|  |  |
| --- | --- |
| **Resolution range** | 57.28 - 2.9 (3.004 - 2.9) |
| **Space group** | P 1 |
| **Unit cell** | 69.1819, 69.2115, 73.1926<br>63.542, 89.304, 60.0427 |
| **Unique reflections** | 22220 (2221) |
| **Completeness (%)** | 98.85 (98.27) |
| **Wilson B-factor** | 71.57 |
| **Reflections used in refinement** | 22190 (2221) |
| **Reflections used for R-free** | 1025 (111) |
| **R-work** | 0.2454 (0.4127) |
| **R-free** | 0.3264 (0.4235) |
| **Number of non-hydrogen atoms** | 6580 |
| - Macromolecules | 6540 |
| - Ligands | 0 |
| - Solvent | 40 |
| **Protein residues** | 808 |
| **RMS (bonds)** | 0.014 |
| **RMS (angles)** | 1.81 |
| **Ramachandran favored (%)** | 89.41 |
| **Ramachandran allowed (%)** | 9.31 |
| **Ramachandran outliers (%)** | 1.28 |
| **Rotamer outliers (%)** | 6.23 |
| **Clashscore** | 21.80 |
| **Average B-factor** | 71.10 |
| - Macromolecules | 69.75 |
| - Solvent | 68.86 |

### Native PAGE

Reactions (10 μL) were set up in Hfq binding buffer (20 mM Tris pH 8.0, 150 mM KCl, 2.5 mM MgCl_2_, 1 mM TCEP) at 30 °C with purified Hfq and 50 nM RNA probes and separated on a 6% (wt/vol) native polyacrylamide gel. The gel was run on TBE buffer for 80 min at 120 V and imaged on a Syngene InGenius3 system. The concentration of Hfq used are indicated on figures.

### RNA sequencing

Bacteria were harvested at N-24. RNA was extracted using the RNAsnap protocol (41). Three biological replicates of each strain were taken and mixed with a phenol:ethanol (1:19) solution at a ratio of 9:1 (culture:solution) before harvesting the bacteria immediately by centrifugation. Pellets were resuspended in RNA extraction solution (18 mM EDTA, 0.025% SDS, 1% 2-mercaptoethanol, 95% formamide) and lysed at 95°C for 10 min. Cell debris was pelleted by centrifugation. RNA was purified with PureLink RNA Mini Kit extraction columns (Invitrogen, 12183018A) and largely in accordance with the manufacturer’s protocol for Total Transcriptome Isolation except with a final ethanol concentration of 66% to increase the yield of smaller RNA species. Analysis of extracted RNA was performed following depletion of ribosomal RNA molecules using a commercial rRNA depletion kit for mixed bacterial samples (Lexogen, RiboCop META, #125). The ribo-depleted RNA samples were first fragmented using ultrasound (4 pulses of 30 s at 4°C). Then, an oligonucleotide adapter was ligated to the 3’ end of the RNA molecules. First-strand cDNA synthesis was performed using M-MLV reverse transcriptase with the 3’ adapter as primer. After purification, the 5’ Illumina TruSeq sequencing adapter was ligated to the 3’ end of the antisense cDNA. The resulting cDNA was PCR-amplified using a high-fidelity DNA polymerase and the barcoded TruSeq-libraries were pooled in approximately equimolar amounts. Sequencing of pooled libraries, spiked with PhiX control library, was performed at a minimum of 7 million reads per sample in single-ended mode with 100 cycles on the NextSeq 2000 platform (Illumina). Demultiplexed FASTQ files were generated with bcl-convert v4.2.4 (Illumina). Raw sequencing reads were subjected to quality and adapter trimming via Cutadapt (42) v2.5 using a cutoff Phred score of 20 and discarding reads without any remaining bases (parameters: --nextseq-trim=20 -m 1 -a AGATCGGAAGAGCACACGTCTGAACTCCAGTCAC). Afterwards, all reads longer than 11 nt were aligned to the *E. coli* K12 MG1655 reference genome (RefSeq assembly accession: GCF_000005845.2) using the pipeline READemption (43) v2.0.3 with segemehl version 0.3.4 (44) and an accuracy cut-off of 95% (parameters: -l 12 -a 95). READemption gene_quanti was applied to quantify aligned reads overlapping genomic features by at least 10 nt (-o 10) on the sense strand (-a) based on RefSeq annotations (CDS, ncRNA, rRNA, tRNA) for assembly GCF_000005845.2 from Mar 11, 2022. Based on these counts, differential expression analysis was conducted via DESeq2 (45) version 1.24.0. Read counts were normalized by DESeq2 and fold-change shrinkage was conducted by setting the parameter betaPrior to TRUE. Differential expression was assumed at adjusted p-value after Benjamini-Hochberg correction (padj) < 0.05 and |log2FoldChange| ≥ 1. DESeq2 analysis can be found in Supplementary Data 2.

## Results

### Characterisation of a systematic alanine mutant library of Hfq

Our objective was to make a systematic alanine mutant library of Hfq to identify amino acids (aa) that are important to the adaptive response to N starvation in *E. coli*. To do so, we first developed a simple experimental system to evaluate the mutant library of Hfq. Bacteria devoid of Hfq (Δ*hfq*) do not display any viability defect under N replate (N+) conditions or following short-term (20 min; N-) N starvation (20). However, the absence of Hfq severely compromises the viability of a population of Δ*hfq E. coli* following 24 h of N starvation (Figure 1B). The proportion of viable bacteria in the Δ*hfq E. coli* population can be restored to wild-type (WT) levels by supplying exogenous Hfq from an inducible plasmid (here L-arabinose was used as the inducer to drive transcription of *hfq* from the *araB* promoter). The plasmid-borne Hfq contained a FLAG-tag at its carboxyl terminal end to allow detection of the protein in bacterial whole-cell extracts (Figure 1C). The Hfq of *E. coli* is 102 aas in length and residues 1-65 represent the conserved core. Given that removal of the CTD aas 73-102 does not compromise Hfq’s function (46), including in its role in the adaptive response to N starvation (Figure 1D), we systematically changed each aa residue between position 3 and 72 to alanine (excluding the 2 alanine residues at position 12 and 56 in the WT Hfq sequence). The mutant Hfq library consisted of 68 mutant proteins.

Plasmids harbouring the Hfq mutants were introduced into Δ*hfq* bacteria, which were then grown under N replete conditions in the presence of L-arabinose. As shown in Figure 1E, the growth dynamics of bacteria expressing the Hfq mutants did not significantly differ from that of WT bacteria. Under our conditions, growth arrest occurs when the sole N source is exhausted and for all strains this occurred ∼3.5 h following inoculation (Figure 1E). We left the cultures under N starvation for 24 h (hereafter referred to as N-24) and enumerated the proportion of viable cells in the population by counting colony forming units. We identified several aa residues in Hfq at which an alanine substitution adversely affected viability of Δ*hfq* bacteria (Figure 2A). For further analyses, we selected Δ*hfq* bacteria encoding plasmid-borne Hfq mutants that displayed a ≥25% reduction (Figure 2A, red line) in the number of viable cells in the population at N-24 compared to Δ*hfq* bacteria encoding plasmid-borne WT Hfq. Twenty five Hfq mutants were selected (Figure 2A, black bars). Of these 25 mutations in Hfq, as shown in Figure 2B, 7 were surface exposed residues at the proximal face (K3, Q8, D9, F39, H57 and Y55; note that K3 is absent from the structure shown in Figure 2B) including the partially exposed residue K56; 5 were surface exposed residues on the distal face (I30, G29, and Y25) including the partially exposed residues L32 and T61; 3 residues lie within the hydrophobic core (V22, G34 and L46) close to the distal face of the Hfq hexamer; two are surface exposed on the rim region (R16 and R17); 7 residues are located at the interface between the individual monomers of the Hfq hexamer (F11, L12, I24, L26, V54, I59 and V62); and 1 residue (I36) was internal to the Hfq monomer. We probed the cell extracts of Δ*hfq* bacteria encoding the 25 plasmid-borne Hfq mutants using anti-FLAG antibodies to determine how the expression levels of the mutant proteins differed from that of the WT protein at N-24. We quantified the intensity of the band corresponding to each mutant protein on the Western blots and considered a mutant to be expressed to near WT level if the band intensity was within ∼75% of the band intensity of WT Hfq. All Hfq mutants, except proximal face mutant Y55A, distal face mutant I30A, the L46A mutant within the hydrophobic core of the Hfq hexamer, several mutations (F11A, L12A, I24A) at the interface between the individual monomers of the Hfq hexamer and the I36A mutant which is internal to the Hfq monomer were expressed at levels comparable to that of the WT Hfq (Figure 2C). Thus, it is conceivable that the aa residues F11, L12, I24, I30, I36, L46 and Y55 contribute to the overall structural integrity and stability of either individual Hfq monomers or the hexamer.

**Figure 2.**
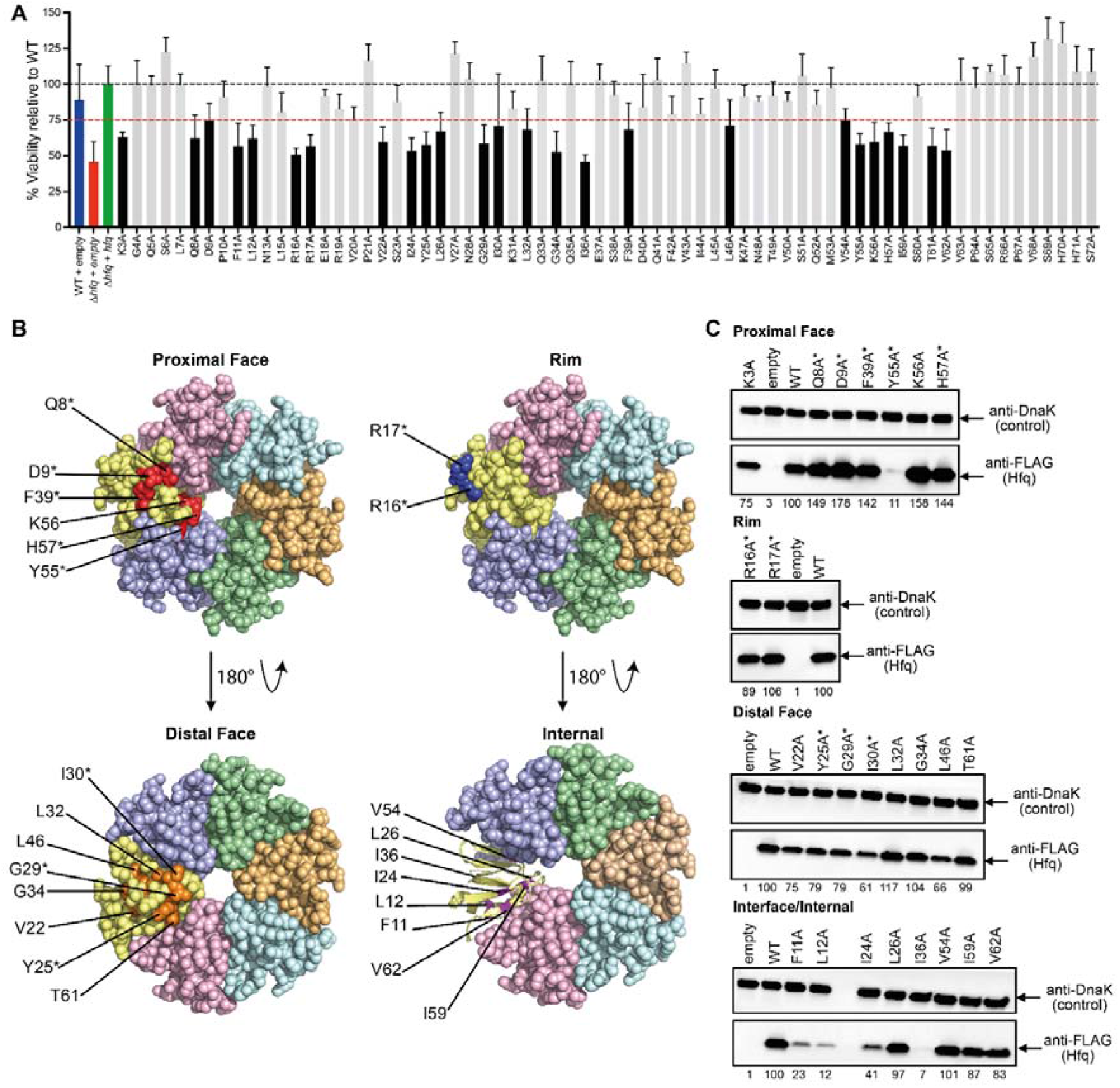
Characterisation of a systematic alanine mutant library of Hfq **(A)** Viability of WT and Δ*hfq E. coli* expressing plasmid-borne alanine mutants of Hfq (pBAD24-*hfq*-3xFLAG) measured by counting CFU following 24 hours of nitrogen starvation (N-24). Values are represented as a percentage of viable cell counts relative to the wild-type complemented strain. The dashed red line indicates ≥25% drop in viable cell count. Error bars represent standard deviation (n = 3). **(B) S**tructures of hexameric Hfq (PDB: 1HK9) with each monomer of Hfq coloured separately. Amino acid residues at which an alanine substitution results in ≥25% decrease in viability with respect to WT (black bars in **(A)**) are labelled and coloured in red, orange, blue and purple for those found at the proximal and distal faces, the rim region and at the monomer-monomer interfaces or internally, respectively (aa K3 is not present in the Hfq structure used here). **(C)** Representative immunoblots of whole-cell extracts of Δ*hfq E. coli* containing either a pBAD18 empty vector control, or WT or alanine mutants of pBAD24-*hfq*-3xFLAG, sampled at N-24. The immunoblots were probed with anti-FLAG antibody and anti-DnaK antibody (loading control). The ratio of the intensity of bands corresponding to alanine mutants of Hfq relative to that of WT Hfq from the same immunoblot are indicated below the immunoblot. Surface exposed residues are indicated by *.

### Molecular dynamics simulations confirm aa residues that contribute to structural integrity of the Hfq monomer

Molecular dynamics (MD) simulation allows the conformational space available to macromolecules to be explored and is an effective method for investigating structural changes induced by mutations. To investigate alanine substitutions at aa residues suspected to be important for the structural integrity of the Hfq monomer, we conducted standard MD simulations on Hfq monomer using two protein force fields (Table 1; see also Materials and Methods for difference between ff12SB and ff19SB protein force fields). We chose Hfq mutants, which appeared to be most structurally destabilised in Figure 2C (F11A, L12A, I24A, I36A and Y55A) for MD analysis (Figure 3A). Briefly, MD plots in Figure 3B, 3D and 3E reveal the percentage of interatomic contacts present in the starting structure (i.e. the native contacts), which are maintained through the course of the MD simulation. Higher populations of fewer native contacts indicate greater departure from the native protein fold. The results revealed visible signs of instability of the native structure in the mutant proteins (Figure 3B). This was especially prominent for the L12A, I24A and I36A mutants (Figure 3B). In the I36A mutant, for example, which is an internal residue in the Hfq monomer, the presence of alanine creates a void for which the protein compensates with structural rearrangements, causing the native fold to split open and fall apart (Figure 3C). However, the structural basis of the disruptive influence of the surface-exposed F11A and Y55A mutation was more elusive. By measuring the root mean square fluctuation (RMSF), which is a measure of fluctuations in a proteins secondary structure during MD simulation, we observed alternative docking arrangements of the α-helix 1 to the rest of the domain and generally increased flexibility in the F11A mutant (Figure 3D). The Y55A mutation led to increased flexibility of the nearby β-sheets as well, eventually causing the β-sheets 4 and 5 to split open (Figure 3D). Overall, all the mutants selected for MD analysis were generally less stable in standard MD simulations than the WT Hfq protein. However, due to limited sampling, we did not observe the loss of the protein structure beyond the early stages of the MD simulations process. Even extending the standard MD simulations up to 20 μs was insufficient to observe complete loss of the characteristic Hfq fold (Figure 3B and Table 1). Therefore, we performed enhanced sampling REST2 simulations, where we could observe virtually complete loss of the native Hfq fold for the mutants selected for analysis (Figure 3E). The only exception was the Y55A mutant which was still relatively stable by the simulation end but still less so compared to the WT protein. Importantly, the WT Hfq protein was always fully stable in both standard MD and REST2 simulations. Since no restraints or artificial biases were at any point applied to reinforce the native structure, the MD analysis conclusively shows that the alanine substitutions at aa residues F11, L12, I24, I36 and Y55 severely disrupt the Hfq monomer structure.

**Figure 3.**
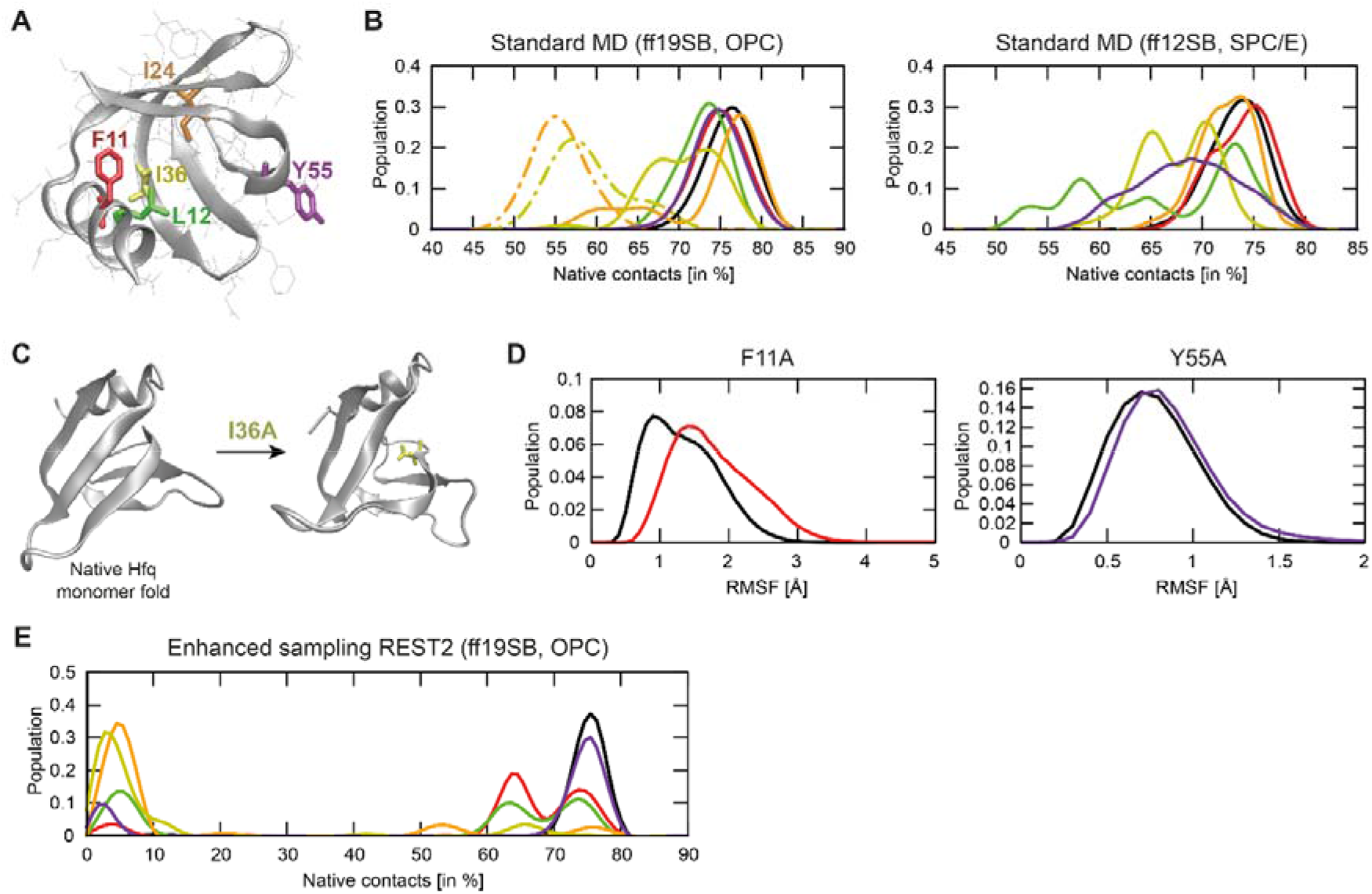
MD simulations of unstable alanine mutants of Hfq **(A)** Structure of the Hfq monomer with the amino acids for which we studied the effects of alanine substitutions using MD simulations; the substitutions are labeled and individually colored. The other amino acid residues are shown as thin grey lines. The secondary structure is indicated with grey ribbon. **(B)** Graphs of the populations of native contacts in standard MD simulations of alanine mutant Hfq protein, colour coded as in **(A)**. The black line corresponds to the WT simulations. The graphs show the percentage of the interatomic contacts present in the starting structure (i.e. the native contacts) that are subsequently maintained in MD simulations of each alanine mutant of Hfq. Higher populations of fewer native contacts indicate more significant departures from the native Hfq fold. The dot-dashed lines indicate the two standard simulations which were extended to 20 μs (see Table 1). For visual clarity, the data curves were smoothed with cubic spline interpolation. **(C)** Collapse of the native Hfq protein fold with I36A mutation as observed by MD simulations. The mutation causes structural collapse of the native fold during which the beta-sheet surface splits open. Very similar changes were also observed with the L12A and I24A mutations. **(D)** Graphs of the average root mean square fluctuations (RMSF) of the α-helix (F11A, red) and β-sheet residues (Y55A, violet) and their comparison with the WT (black curve) in standard MD simulations using ff19SB. **(E)** Graph as in **(B)** but of the native contacts in the reference replica of the REST2 enhanced sampling MD simulations.

Lastly, we note both the ff12SB and ff19SB protein force fields revealed only minor differences within the limit of the sampling and the same qualitative trends in terms of stability of the mutants (Figure 3B). The exception seems to be the Y55A mutant which was significantly less stable with ff12SB than with ff19SB (Figure 3B). In general, the ff19SB maintained slightly more native contacts but the observed differences are very minor. The specific water models (OPC vs SPC/E) utilized with each force field could also be the source of these differences.

### Alanine substitutions at conserved residue V22 or G34 do not alter structural integrity or RNA binding activity of Hfq in vitro

Of the Hfq mutants that displayed compromised viability at N-24 but were expressed to near or at wild-type levels (18 in total), several (13) have been previously implicated to be important for or involved in RNA binding by Hfq (46–48). These are K3, Q8, D9, R16, R17, Y25, G29, L32, F39, K56, I59, H57 and T61 (Figure 1A and Figure 2A). To the best of our knowledge, residues V22, G34, L26, V54 and V62 have not been previously implicated in Hfq function. V22 and G34 are within the hydrophobic core of Hfq monomer close to the distal face and residues L26, V54 and V62 are at the monomer-monomer interface. Alanine substitutions at the monomer-monomer interface could affect the conformation of Hfq hexamer, rendering it functionally ineffective during N starvation. As residues V22 and G34 are fully conserved in Hfq from different bacteria (Supplementary Figure 1) and located in the hydrophobic core of Hfq, close to the distal face, we focused our analysis on investigating why alanine substitution at these positions compromised Hfq function under N starvation. Standard MD simulation of both V22A and G34A Hfq mutants revealed no signs of instability of the native structure and similar number of native contacts were maintained by both mutants as in the WT Hfq monomer (Figure 4A). However, we note subtle differences in the native contacts maintained between the G34A and the WT Hfq monomer (Figure 4A). Consistent with the MD simulation data, CD spectroscopy analysis in the UV range of V22A and G34A Hfq mutants revealed no changes between samples at 25°C and 95°C, indicating that the stability of the secondary structure of the mutants did not substantially differ from that of the WT Hfq protein (Figure 4B). However, we do note small differences when comparing the spectra of each protein (Figure 4B), possibly indicating subtle changes in the secondary structure (which are nonetheless stable) between the mutant and WT Hfq. Further, in the near-UV range, we observed consistent changes in the spectra between 25°C and 95°C between each protein, indicating a comparable level of tertiary structure stability (Figure 4C). Nonetheless, at 25°C the near-UV spectra of both mutants are similar and characteristic of that of the WT protein (Figure 4C). Collectively, the CD spectroscopy analysis shows that alanine substitution at either position V22 or G34 are unlikely to cause large scale structural perturbations or instability in Hfq.

**Figure 4.**
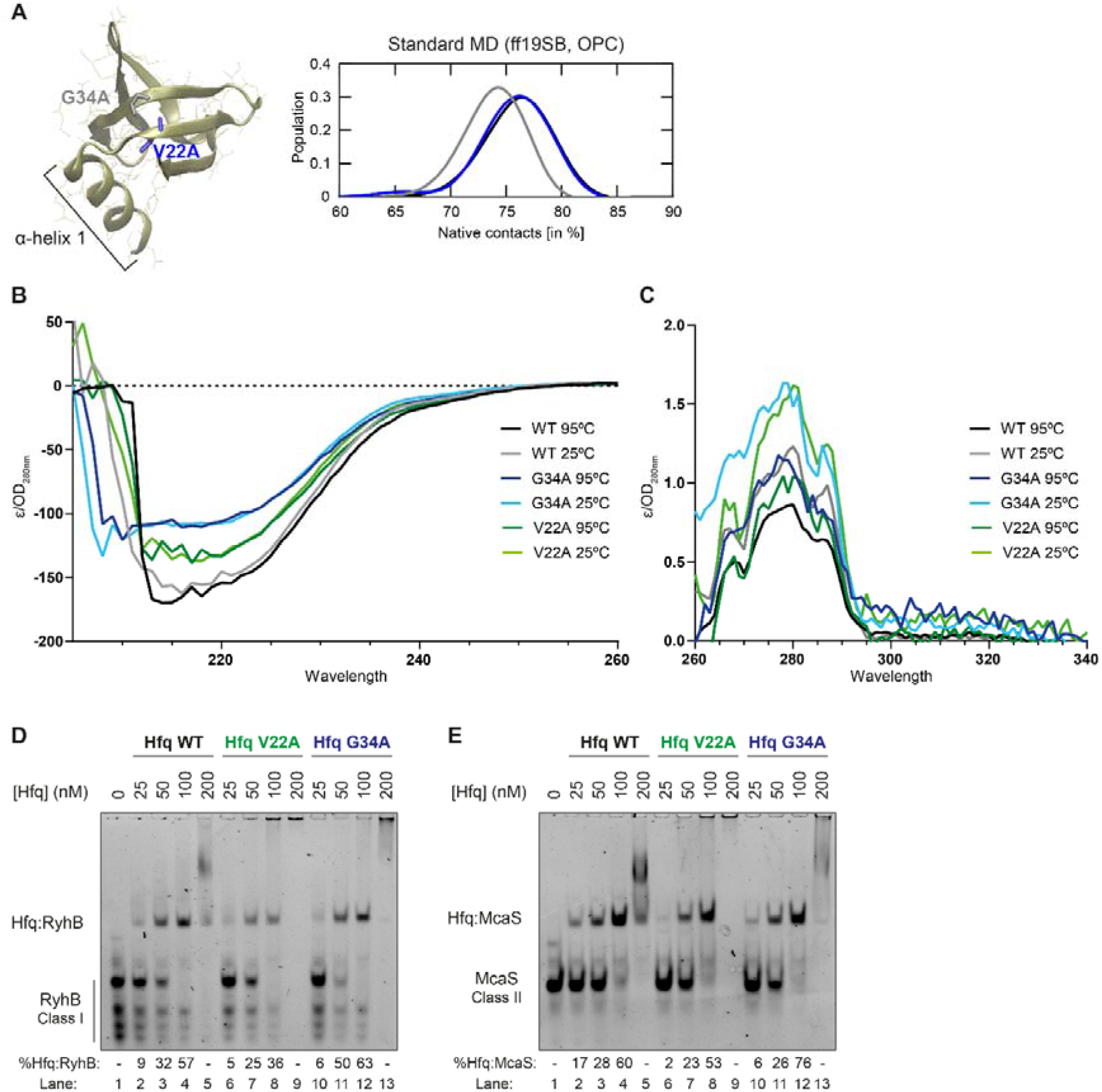
Properties of V22A and G34A alanine mutants of Hfq. **(A)** Structure of the Hfq monomer indicating amino acid residues V22 and G34; α-helix 1 is indicated. Graphs of the native contacts in standard MD simulations (ff19SB, OPC) of alanine mutant Hfq protein (as in Figure 3B). The black line corresponds to the WT simulations, with the blue and grey lines corresponding to the V22A and G34A Hfq alanine mutants, respectively. The graphs indicate how many of the interatomic contacts present in the experimental WT structure (i.e. the native contacts) are subsequently maintained in MD simulations of each system. Higher populations of fewer native contacts indicate more serious departures from the native Hfq fold. **(B)** Circular dichroism analysis of WT, V22A and G34A alanine mutants of Hfq protein, at 25°C and 95°C between wavelengths of 205- 260nm. **(C)** as in **(B)**, but between wavelengths of 260-340nm. **(D)** Representative native-gel showing WT, V22A and G34A alanine mutants of Hfq proteins binding to RhyB sRNA. Bands corresponding to free (unbound) RyhB and RyhB in complex with Hfq (Hfq:RyhB) are indicated. The percentage of the total signal within each lane which corresponds to the Hfq:RyhB complex is indicated; ‘-‘ indicates that the Hfq:RyhB complex was either undetectable or less than 1% of the total lane signal. We note some degradation of the RyhB probe, resulting in multiple extra bands. **(E)** As in **(D)** but with McaS sRNA.

Next, we considered whether the V22A and G34A Hfq mutants are defective for sRNA binding. To investigate this, we conducted electrophoretic mobility shift assays with purified proteins and representative class I (RyhB) and class II (McaS) sRNA probes. At equimolar concentrations of Hfq and either RyhB (Figure 4D) or McaS (Figure 4E) probe, the binding of both mutant Hfq proteins to the sRNA probes was comparable to that of the WT protein (lanes 3, 7 and 11). However, we noted a tendency of the V22A mutant to aggregate at concentrations >100 nM, potentially explaining the reduced intensity of the band corresponding to the V22A Hfq:RyhB/McaS complexes compared to those of the G34A or WT Hfq protein. Overall, it seems that the presence of an alanine residue at either position V22 or G34 does not markedly alter the ability of Hfq to interact with sRNA *in vitro*, regardless of the mode of interaction of the sRNA with Hfq, under equilibrium conditions. Further, the electrophoretic properties (i.e. size, shape and location on the gel) of RyhB and McaS complexes with either of the Hfq mutants are comparable to those of WT Hfq, suggesting that that the overall structural conformation and the ability of the V22A or G34A Hfq mutants to hexamerise is not compromised, consistent with the MD simulation of the monomers of Hfq mutants and CD spectroscopy analysis of purified proteins.

To further elaborate on the properties of V22A Hfq mutant, we obtained diffracting co-crystals of the V22A mutant using poly(A) RNA (pA_4_) RNA. The crystals diffract to roughly 3.0 angstroms, but with anisotropy, and the structure was solved using molecular replacement of the core residues 6-66 of *E.coli* Hfq, based on PDB entry 3GIB (12). Two Hfq hexamers occupy the asymmetric unit, leaving 38% solvent content, and the hexamers stack tightly through interactions of the distal face and circumferential rim, but leave gaps between the stacks on the proximal face (Supplementary Figure 2). The space in the lattice is likely filled with the N- and C-terminal tails of Hfq, but these are comparatively disordered and could not be modelled in the crystal. The gap spacing is likely maintained through weak, multivalent interactions and may be a balance of attractive and electrostatically repulsive interactions, in analogy to liquid-liquid phase separations. Nonetheless, the diffraction intensities are anisotropic, which may reflect the disorder due to imperfect packing in those proximal-proximal interfacial packing gaps. On the proximal side, we observe density which could fit adenines, which would represent a non-canonical interaction with pA_4_ RNA on this surface (Supplementary Figure 2). However, the electron density is not satisfactory to model the details of these interactions, and the result must be interpreted with caution at this point.

One molecule of pA_4_ RNA could be traced at the distal face of one of the hexamers, and disordered portions of the RNA polymer could be fit at the distal surface elsewhere on the hexamer (Figure 5A and Supplementary Figure 2). An overlay of the WT and V22A Hfq mutant crystal structures reveal similar conformations around the site of substitution, within the error of the intermediate resolution of the crystal structure of the mutant protein (Figure 5B). The so-called A-R-N motif (where A is adenosine, R a purine and N any nucleotide), found upstream regions of those mRNAs that are regulated by sRNAs, specifically interacts with the distal RNA binding face of Hfq. Focussing on the interaction of the distal face with the pA_4_ RNA, we found that the interactions of the V22A Hfq mutant are very close to those seen for the canonical A-R-N pattern recognition that has been well characterised by earlier crystallographic studies for the WT protein (Figure 5C). We conclude that, the alanine substitution at conserved aa V22 only subtly, if at all, alters the structural confirmation and the mode of RNA interaction of the mutant protein. Overall, the independent *in vitro* assays reveals that an alanine substitution at the conserved residue V22 or G34 do not alter the structural or RNA binding activity of the mutant proteins.

**Figure 5.**
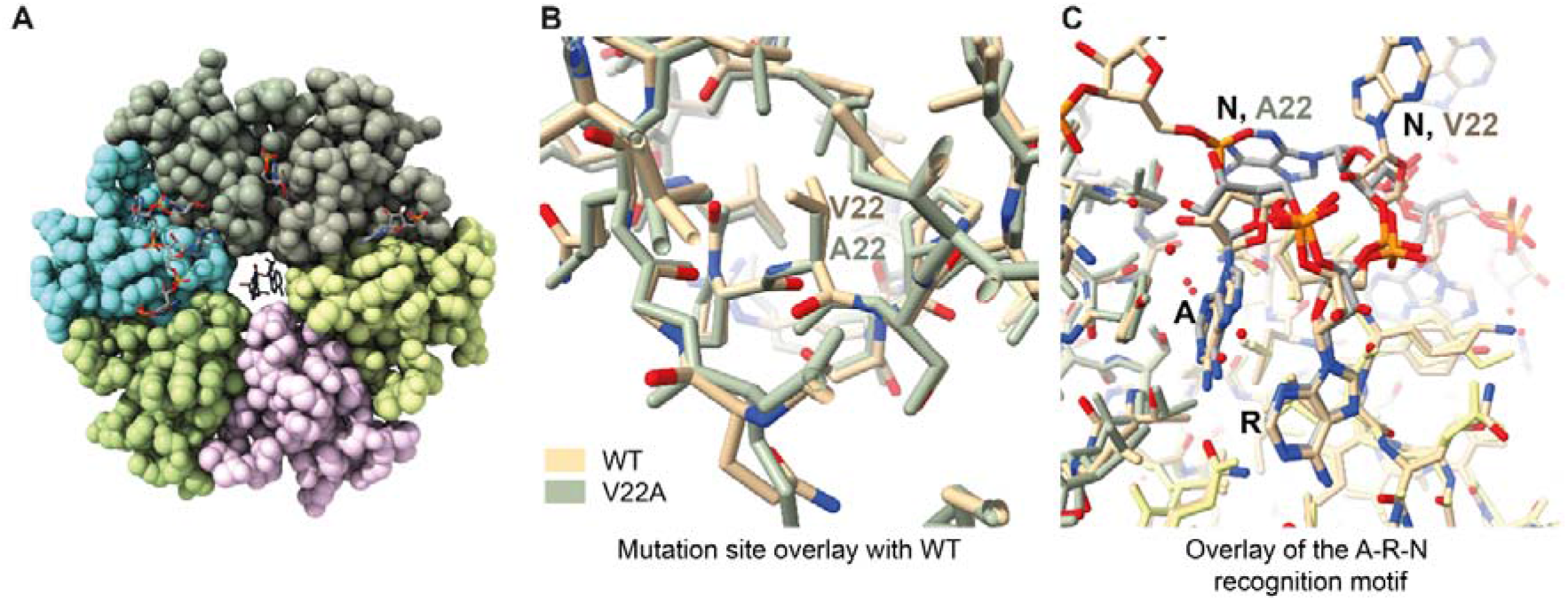
Crystal structure of *E. coli* Hfq V22A in complex with pA_4_ **(A)** 3D structure of V22A Hfq, with one pA_4_ molecule making canonical A-R-N interactions. Adenines are seen with partial occupancy at the other side of the distal face, but the lattice contacts may disfavour their formation. The hole in the centre reveals two adenines that are interacting with the proximal face in a non-canonical interaction mode. **(B)** Overlay of the mutant and wild type subunit with zoom onto the site of mutation, revealing little conformational change. Wild-type structure is shown in cream, and the V22A mutant structure is shown in green-grey. **(C)** Overlay of the mutant and wild-type subunit showing the similarity of the interaction with the A-R-N recognition motif. The N nucleotide differs in conformation, most likely as a result of the crystal lattice packing.

### The transcriptomes of V22A and G34A mutants resemble that of bacteria devoid of Hfq

To understand why bacteria expressing the V22A or G34A Hfq mutants are compromised for surviving long-term N starvation, despite being structurally intact and able to interact with both classes of sRNAs, we compared the transcriptomes of bacteria expressing WT Hfq with that of bacteria containing the V22A and G34A Hfq mutants at N-24. Surface exposed residue Q8, at the proximal face of Hfq, is important for Hfq-mediated post-transcriptional regulation involving both class I and class II sRNAs. Thus, we also obtained the transcriptomes of N-24 bacteria expressing the Q8A Hfq mutant and bacteria devoid of Hfq (Δ*hfq*) as positive controls to evaluate the extent of perturbation in the transcriptome of N-24 bacteria containing the V22A and G34A Hfq mutants. We defined differentially expressed genes as those with expression levels changed ≥2-fold with a false discovery rate adjusted P<0.05. As shown in Figure 6A-D, the expression of 306 and 376 genes were upregulated, and 357 and 413 genes were downregulated in bacteria expressing V22A and G34A alanine mutants of Hfq respectively (Figure 6A-D), comparable to the number dysregulated in bacteria devoid of *hfq* (360 up, and 420 down). Put differently, the extent of perturbation to the transcriptome in bacteria encoding the V22A or G34A Hfq mutant is comparable to that in the transcriptomes of bacteria expressing the Q8A Hfq mutant or entirely devoid of Hfq (Δ*hfq*). In further support of this view, we next compared the degree to which genes were differentially expressed between the different strains. Correlation analysis of the fold change of the differentially expressed genes in the transcriptomes of bacteria containing the Q8A, V22A, G34A and Δ*hfq* with each other revealed that the direction in which the transcriptomes are perturbed did not substantially differ (Figure 7). Put simply, broadly the same genes were differentially regulated in the transcriptomes of all bacteria regardless of which aa residue was mutated and to a similar extent as in the transcriptome of Δ*hfq* bacteria. It thus seems that an alanine substitution at V22 or G34 in Hfq has a comparable impact on gene expression during N starvation as in bacteria encoding the Q8A Hfq mutation or devoid of Hfq.

**Figure 6.**
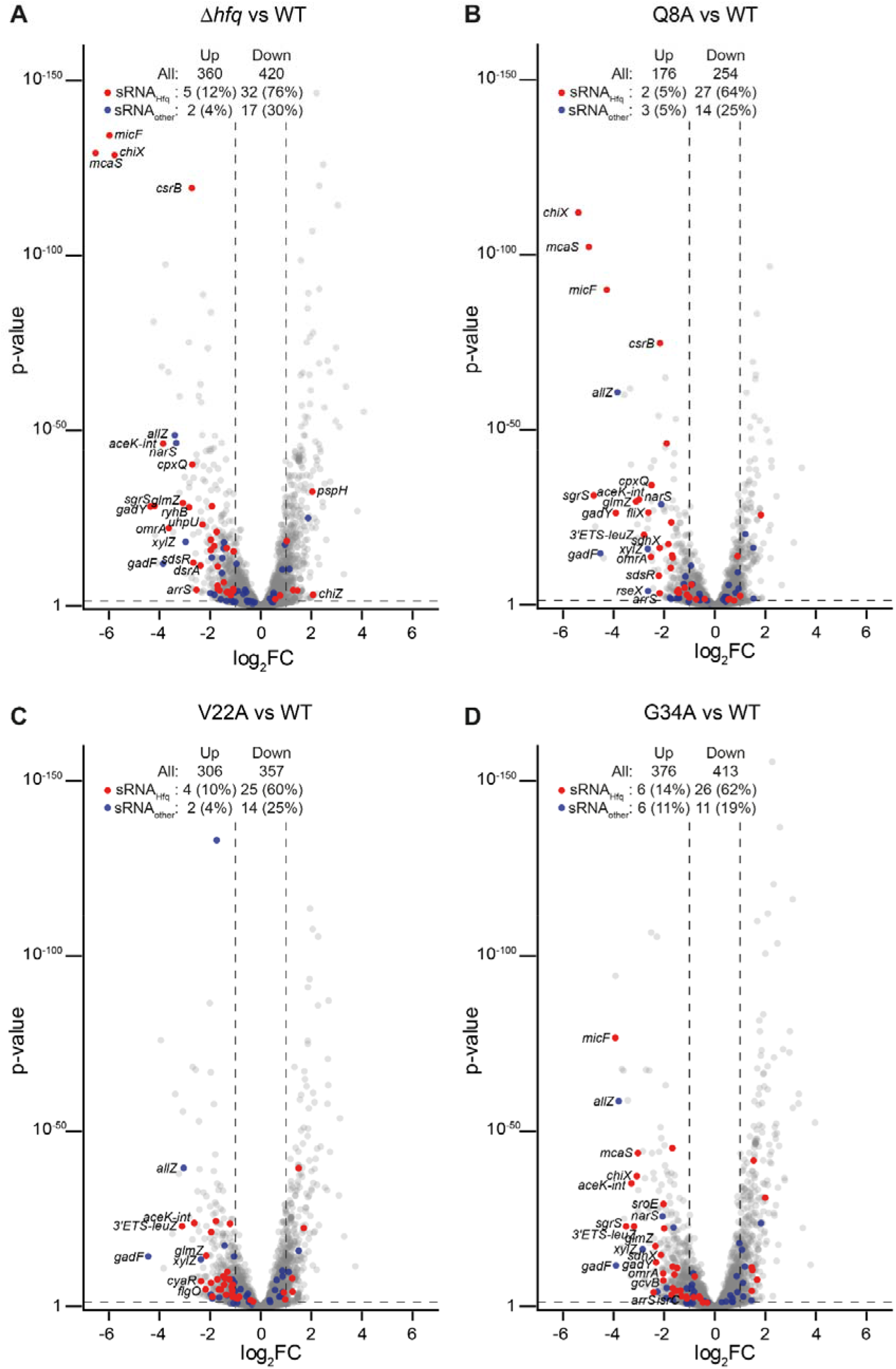
Alanine substitution at Hfq aa residues V22 or G34 lead to substantial changes in the transcriptome. (A) Volcano plots of differential RNA abundance in the transcriptome of Δ*hfq E. coli* versus Δ*hfq E. coli* expressing plasmid-borne WT Hfq. **(B)** As in **(A)** but Δ*hfq E. coli* expressing plasmid-borne Q8A Hfq versus Δ*hfq E. coli* expressing plasmid-borne WT Hfq. **(C)** As in **(A)** but Δ*hfq E. coli* expressing plasmid-borne V22A Hfq versus Δ*hfq E. coli* expressing plasmid-borne WT Hfq. **(D)** As in **(A)** but Δ*hfq E. coli* expressing plasmid-borne G34A Hfq versus Δ*hfq E. coli* expressing plasmid-borne WT Hfq. Analysis was performed by DESeq2. sRNA which have been previously shown to interact with Hfq during N starvation conditions (sRNA_Hfq_) are shown in red, and other sRNA and ncRNA (sRNA_other_) are shown in blue. sRNA and ncRNA differentially expressed more than 2 log_2_ (i.e., a greater than 4-fold change) are labelled. The number and percentage (of total detected) of differentially expressed genes, sRNA_Hfq_ and sRNA_other_ are indicated (log_2_FC > 2 and p-value <0.05).

**Figure 7.**
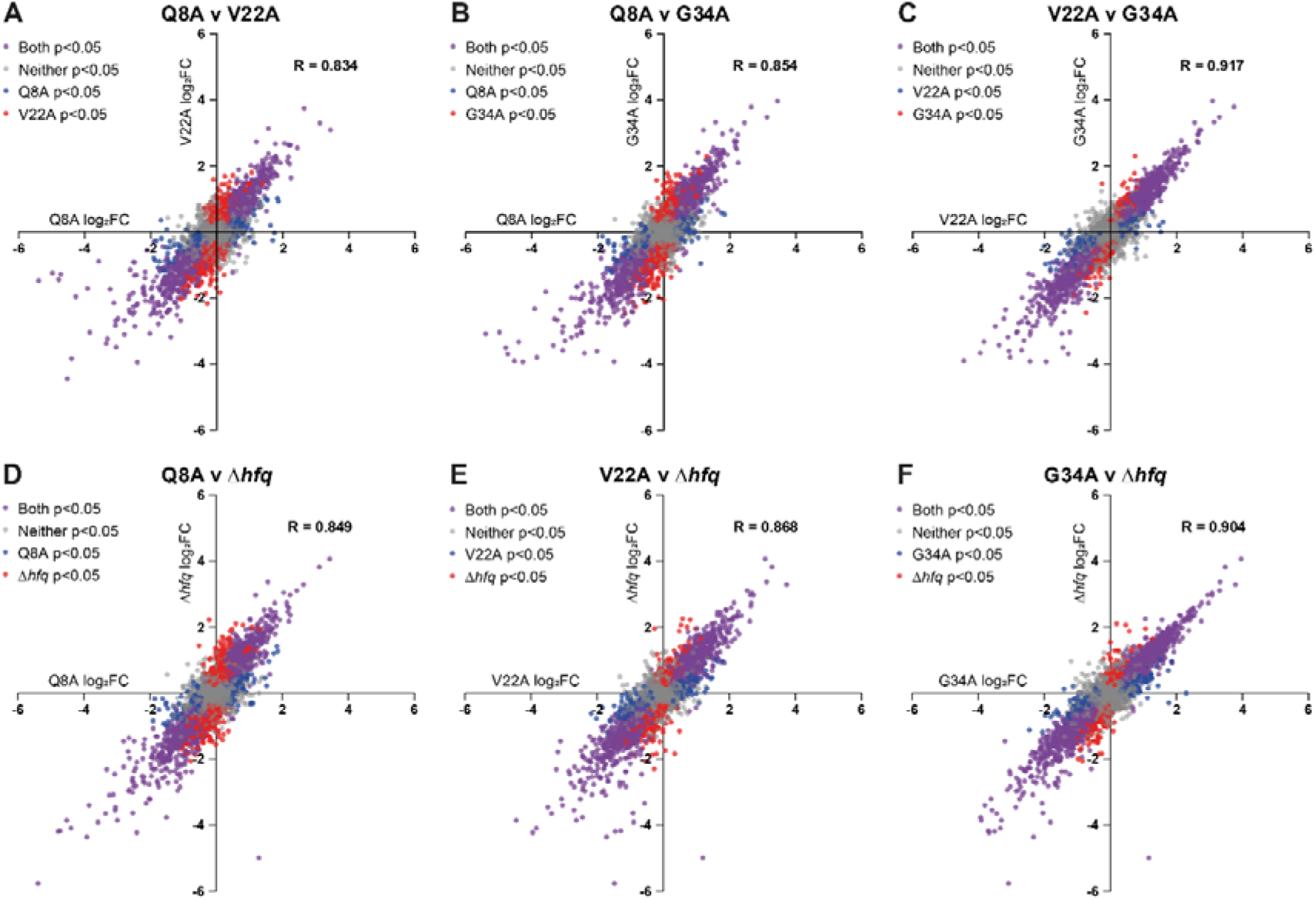
Perturbations to the transcriptome of bacteria expressing alanine mutants of Hfq largely correlate. Differential gene expression, relative to wild-type, was compared between bacteria either lacking Hfq, or containing different mutants of Hfq. Scatterplots comparing log_2_FC of individual genes in Δ*hfq E. coli* or Δ*hfq E. coli* expressing plasmid- borne Q8A, V22A & G34A alanine mutants of Hfq (log_2_FC relative to Δ*hfq E. coli* expressing plasmid-borne WT Hfq) is shown. Genes found to be differentially expressed with respect to bacteria expressing plasmid-borne WT Hfq with a p-value < 0.05 in both pairwise comparisons are shown in purple; genes found to be differentially expressed relative to bacteria expressing plasmid-borne WT Hfq in just one of the compared strains are indicated in blue or red (see inset key for each scatterplot); genes found not to be differentially regulated relative to bacteria expressing plasmid-borne WT Hfq are shown in grey. Pearson correlation coefficient (R) for each comparison is shown.

### The transcriptomes of V22A and G34A mutants reveal destabilisation of sRNA indiscriminately of their mode of interaction with Hfq

As Hfq mediates most of its regulatory function by stabilising sRNAs and facilitating their binding to their cognate target RNA transcripts, we focused on the expression levels of sRNAs in the transcriptomes of bacteria encoding the V22A and G34 to better understand their role in Hfq function. From the literature, we expected that both class I and class II sRNAs to be destabilised in the transcriptome of the Q8A mutant (recall that the proximal face of Hfq is required for binding to both classes of sRNAs). However, from the results of the electrophoretic gel mobility assay shown in Figure 4D and Figure 4E, we did not expect many sRNAs to be destabilised in the transcriptomes of the V22A and G34A mutants. Surprisingly, as shown in Figure 6A-D (red and blue dots) both classes of sRNA transcripts, particularly those that interact with Hfq during N starvation ((19); Figure 6A-D, red dots), were downregulated in all the transcriptomes, consistent with the fact that Hfq is required for sRNA stability. Put differently, we noted that an equal number of sRNA transcripts were downregulated in bacteria containing the V22A or G34A Hfq mutant as in bacteria containing the Q8A Hfq mutant, which is known to be compromised for both classes of sRNA binding *in vitro* (17). We did not observe any correlation between how the sRNAs are made, or whether they are further processed following transcription, with the degree of their destabilisation (Figure 8A). However, the extent by which some of the sRNA transcripts were downregulated substantially differed. For example, the sRNA ChiX is downregulated by ∼42- and ∼54-fold in the bacteria containing the Q8A mutation or devoid of Hfq, respectively, but only by ∼3 and ∼8-fold in bacteria containing the V22A and G34A mutation, respectively (Figure 8A). We note that the majority of sRNA that are known to interact with Hfq during N starvation (19) were destabilised in the context of the Q8A, V22A or G34A mutants. However, this was not the case for non-coding RNA transcripts (which includes sRNAs) that do not interact with Hfq during N starvation (19) (Figure 8B), underscoring the pivotal role Hfq has in sRNA stabilisation. In sum, the results reveal that an alanine substitution at residues V22 or G34 in the hydrophobic core of Hfq, close to the distal face, lead to the destabilisation of sRNAs indiscriminate of their mode of interaction with Hfq.

**Figure 8.**
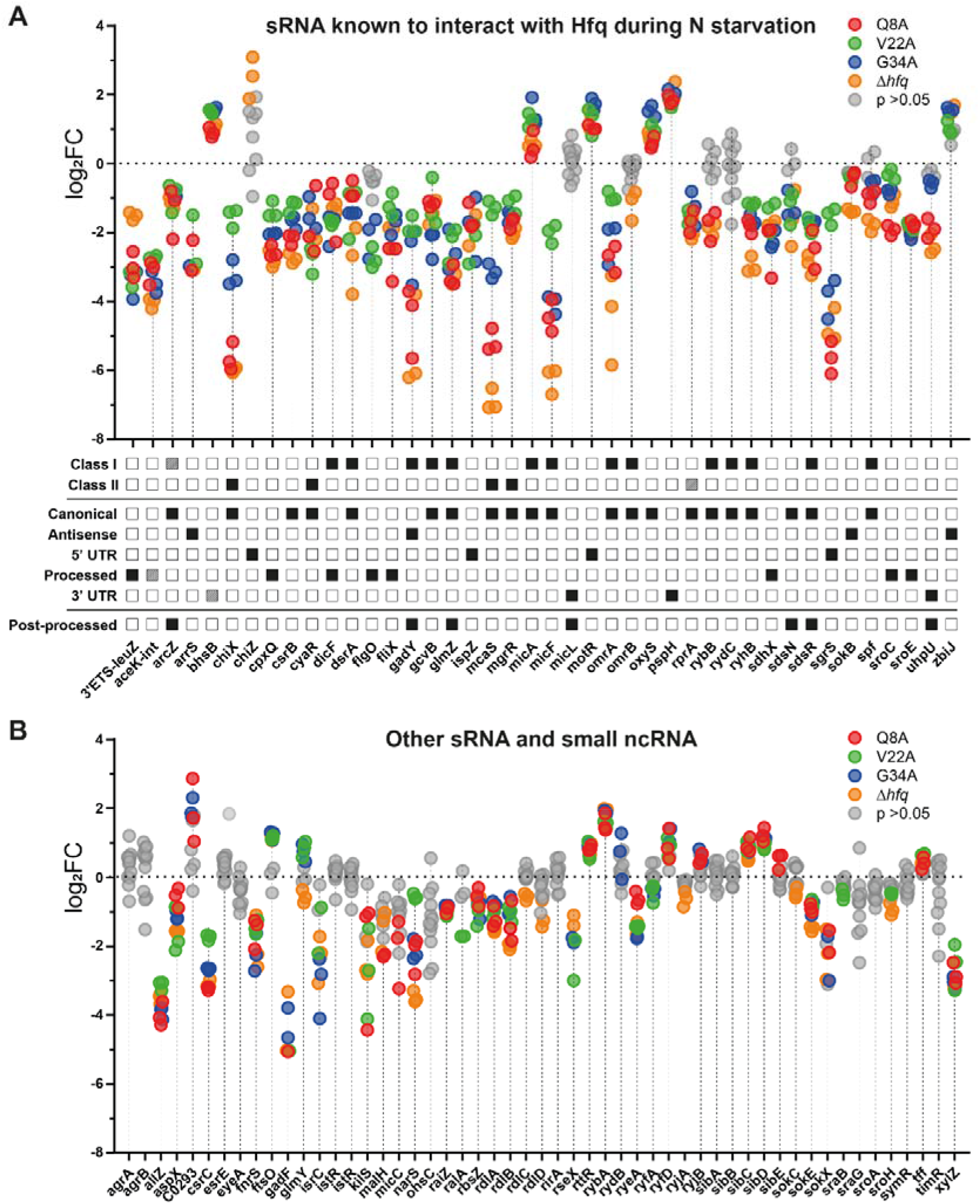
Alanine substitution at Hfq aa residues V22 or G34 lead to indiscriminate destabilisation of Hfq-associated sRNA. (A) Dot plot showing log_2_FC of individual sRNA previously shown to interact with Hfq during N starvation in Δ*hfq E. coli* and Δ*hfq E. coli* expressing plasmid-borne Q8A, V22A & G34A alanine mutants of Hfq, all relative to Δ*hfq E. coli* expressing plasmid-borne WT Hfq. Results from strains containing Q8A, V22A and G34A Hfq and from the Δ*hfq* strain are shown in red, green, blue and orange respectively. Results where an sRNA was not found to be differentially expressed with a p-value <0.05 by DESeq2 in a given strain are shown in grey. The sRNA class (where experimentally established), the nature of their biogenesis, and whether the sRNA undergoes processing following biogenesis is shown below the dot plot. Dashed squares indicate cases where there is uncertainty whether the given sRNA displays the property listed. **(B)** As in **(A)** but for sRNA and ncRNA not detected to interact with Hfq during N starvation.

## Discussion

Hfq is central to the post-transcriptional regulation of gene expression in diverse bacterial species. Much of our understanding of structure-function relationships in Hfq has come from reductionist *in vitro* analyses involving a limited number of prototypical sRNAs and their cognate RNA partners. However, emerging data from genome-wide mapping of the RNA species that interact with Hfq *in vivo* (2,49–53), have revealed that Hfq can facilitate several hundred RNA-RNA interactions, with the complexity of Hfq-mediated RNA-RNA interaction networks compounded by differing expression levels and binding affinities of different RNA species for each other and Hfq. Our current understanding of how Hfq interacts with RNA is based on many *in vitro* biochemical, biophysical and structural studies (e.g.: (10,17,18,46,48,54,55)). These have revealed that surface exposed conserved aa residues on three faces on the toroidal structure of the Hfq hexamer (proximal, distal and rim) cooperatively contribute to bind different species of RNA. However, whilst these *in vitro* studies have provided important mechanistic details into Hfq-mediated regulation of RNA they do not capture the inherent complexity of the cellular system. Hence, as highlighted by the current study, some molecular details of Hfq-dependent regulation of gene expression remained elusive. We have revealed that conserved residues V22 and G34 within the hydrophobic core, close to the distal face of the Hfq hexamer, have a profound impact on the RNA binding activity of Hfq. Alanine substitution at V22 or G34 neither adversely impact the ability of mutant proteins to bind RNA in an equilibrium binding assay *in vitro* nor the overall structural integrity of the protein. Further, the 3D structure of the V22A mutant of Hfq was found to be largely unchanged relative to the native protein (Figure 5). However, strikingly, bacteria expressing either the V22A or G34A Hfq mutant display transcriptome-wide destabilisation of sRNAs. This, consequently, leads to global dysregulation of gene expression, resembling that seen in bacteria devoid of Hfq. Notably, both classes of sRNAs are indiscriminately destabilised in bacteria expressing either the V22A or G34A Hfq mutant, suggesting that alanine substitution at either V22 or G34 adversely impacts both the proximal (impacting both class I and class II) and distal (impacting class II sRNAs) RNA binding faces of Hfq. Indeed, V22 and G34 pack into a hydrophobic pocket and an alanine substitution could impact on the presentation of the α-helix 1 containing Q8 (Figure 4A), which contributes to both class I and class II sRNA interaction at the proximal face of Hfq. We propose that this hydrophobic pocket could be targeted for the rational design of compounds to interfere with the RNA binding activity of Hfq.

In previous work, we reported that the relative sRNA abundance increases over time under N starvation, whereas Hfq levels remain relatively consistent (20,21,56). The Hfq-mediated RNA-RNA interaction network, under a given condition, likely exists in an equilibrium state where individual Hfq-RNA-RNA interactions are in a constant state of flux depending on the transcript levels and relative binding affinities of the different RNA species, with the equilibrium state shifting when the abundance of sRNA changes. This equilibrium of Hfq-RNA-RNA interactions likely has an important role in maintaining homeostasis in the post-transcriptional control of genetic information flow, to a degree we still do not fully understand. We propose that any subtle changes inflicted on RNA binding kinetics resulting from the V22A or G34A mutations, which are difficult to capture with *in vitro* equilibrium binding assays (Figure 4D and Figure 4E), become exacerbated in the cellular context where competition for Hfq occupancy is high. This, consequently, could compromise the equilibrium of Hfq-RNA-RNA interactions and lead to large-scale perturbation of gene expression, akin to that seen in bacteria devoid of Hfq.

Hfq’s contribution to regulating bacterial gene expression extends beyond facilitating RNA-RNA interactions or conferring sRNA stability, and include the biogenesis of ribosomes (57) and tRNA modification (58). Therefore, perturbations to Hfq function will have a direct impact on the cellular proteome. Indeed, we note that the proteome of N-24 bacteria devoid Hfq is substantially perturbed compared to that of WT bacteria (Supplementary Figure 3). Andrade et al (57) reported that mutations in the distal face of Hfq adversely impacted maturation of 16S ribosomal RNA. It is thus possible that subtle conformational changes to the hydrophobic pocket near the distal face of Hfq into which V22 and G34 pack into, also have an adverse effect on 16S ribosomal RNA maturation and, subsequently, on the assembly of 30S ribosomal subunit, impacting cellular translation during long-term N starvation. The study of how individual mutations in Hfq impacts the proteome will be the subject of future research.

In sum, this study underscores the value of using systems level analysis to probe the structure-function relationships of global, nucleic-acid-binding regulators of gene expression, such as Hfq. We have shown that subtle changes in such a protein’s nucleic acid binding activities, which might elude *in vitro* analyses, become amplified in the cellular context providing novel insights into the protein’s functions. Such analyses thus have the potential to identify unknown Achilles’s heels within nucleic acid binding proteins for the rational design of, in this case, antibacterial compounds.

## Acknowledgements

We thank Elena Katzowitzsch from the Core Unit Systems Medicine at the University of Würzburg for excellent technical support for the RNA-seq data generation and analysis.

## Data availability

The RNA-seq discussed in this publication is accessible through ArrayExpress: E-MTAB- 14054. The proteomics data discussed in this publication can be accessed through PRIDE: PXD045656.

---

**Supplementary Figure 1.**
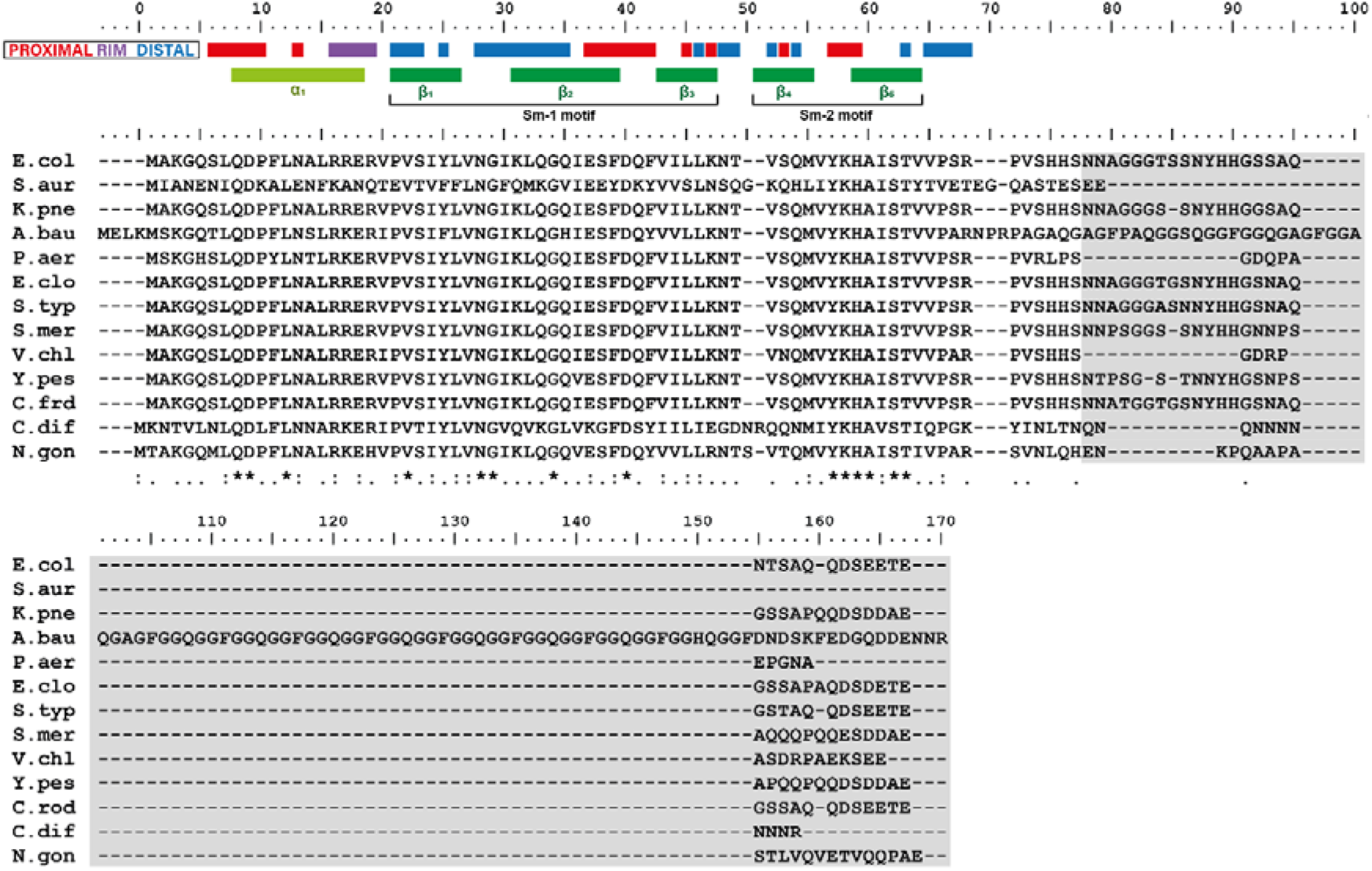
Alignment of Hfq amongst clinically relevant bacterial species. Fully, partially, and poorly conserved residues are indicated below the alignment with * : and . respectively. Residues found at the proximal, rim and distal RNA-binding faces are indicated with red, purple and blue respectively. α-helix, β-sheets, and Sm-like fold motifs are also indicated. Residues highlighted in grey correspond to the disordered C-terminal tail. Abbreviations correspond to the following species: E.col, Escherichia coli K12-MG1655; S.aur, Staphylococcus aureus NCTC 8325; K.pne, Klebsiella pneumoniae HS11286; A.bau, Acinetobacter baumannii ATCC 19606; P.aer, Pseudomonas aeruginosa PAO1; E.clo, Enterobacter cloacae ATCC 13047; S.typ, Salmonella typhi CT18; S.mer, Serratia marcescens Db11; V.chl, Vibrio cholera 10432-62; Y.pes, Yersinia pestis D182038; C.frd, Citrobacter freundii CFNIH01; C.dif, Clostridioides difficile 630; N.gon, Neisseria gonorrhoeae FA1090.

**Supplementary Figure 2.**
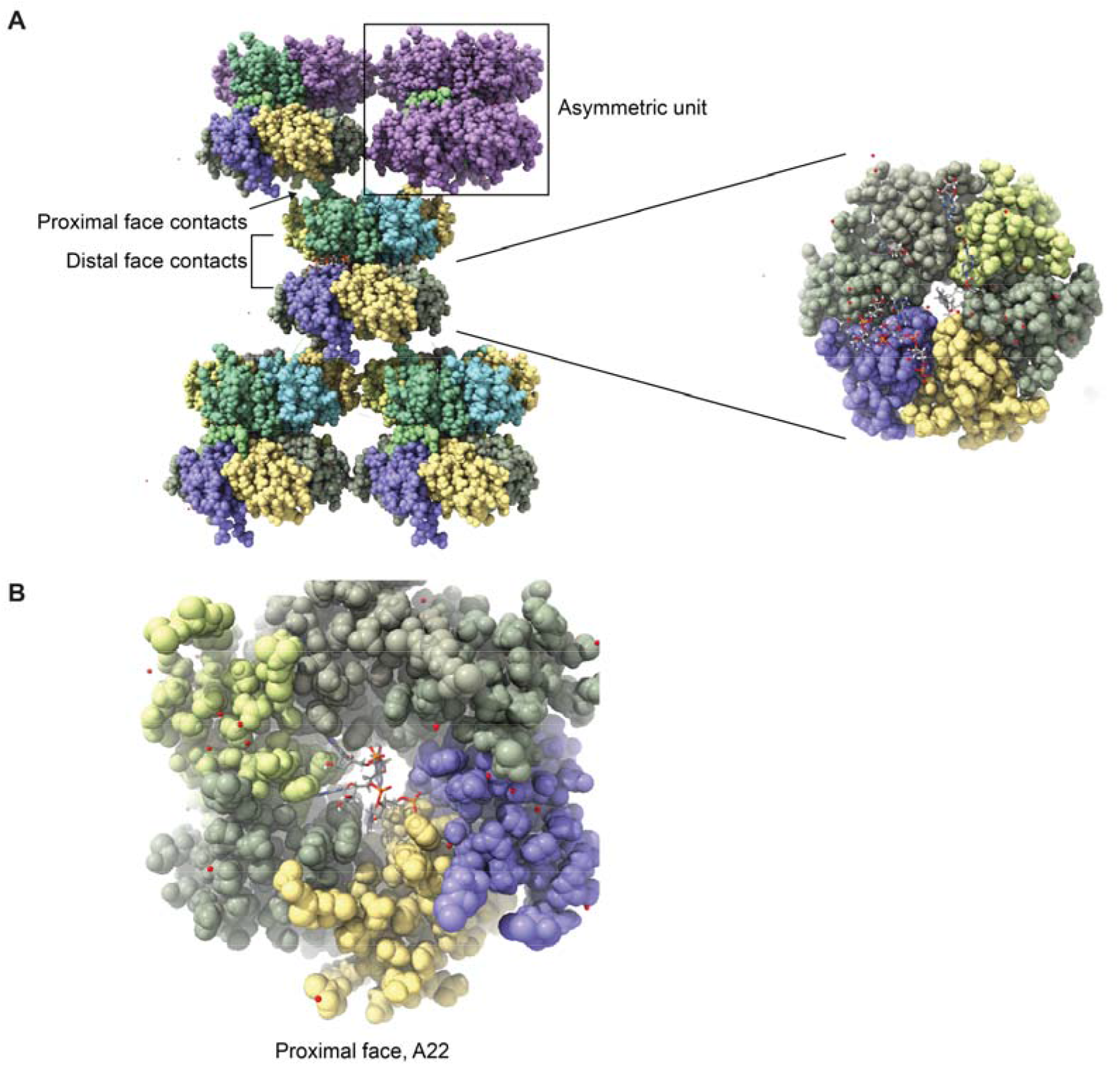
(A) The crystal lattice organisation of V22A Hfq pA_4_. Two hexamers of Hfq occupy the asymmetric unit. The two hexamers stack through interactions of the distal face, and in the perpendicular directions, through the circumferential rim. The distal interfacial contacts include shared interactions with RNA, and the panel on the right shows the view into the distal face of the lower of the two hexamers, with one pA_4_ molecule making canonical A-R-N interactions. Adenines are seen with partial occupancy at other sites of the distal face, but the lattice contacts may disfavour their formation. The hole in the centre reveals an adenine that is interacting with the proximal face in a non-canonical interaction mode. In contrast to the distal face packing, the proximal face has a gap that is occupied by disordered regions of the Hfq termini (typically 1-5 and 67 to 102). These appear to form a densely packing interface with high degree of disorder and might resemble the ensemble conformations associated with liquid phase separation state. **(B)** The interactions of A with the proximal surface. The electron density, which is not well resolved, has approximate three fold symmetry and is likely to represent an average of six orientations of the base. Other adenine bases may partially occupy part of the pockets that are preferred binding sites for uracil bases.

**Supplementary Figure 3.**
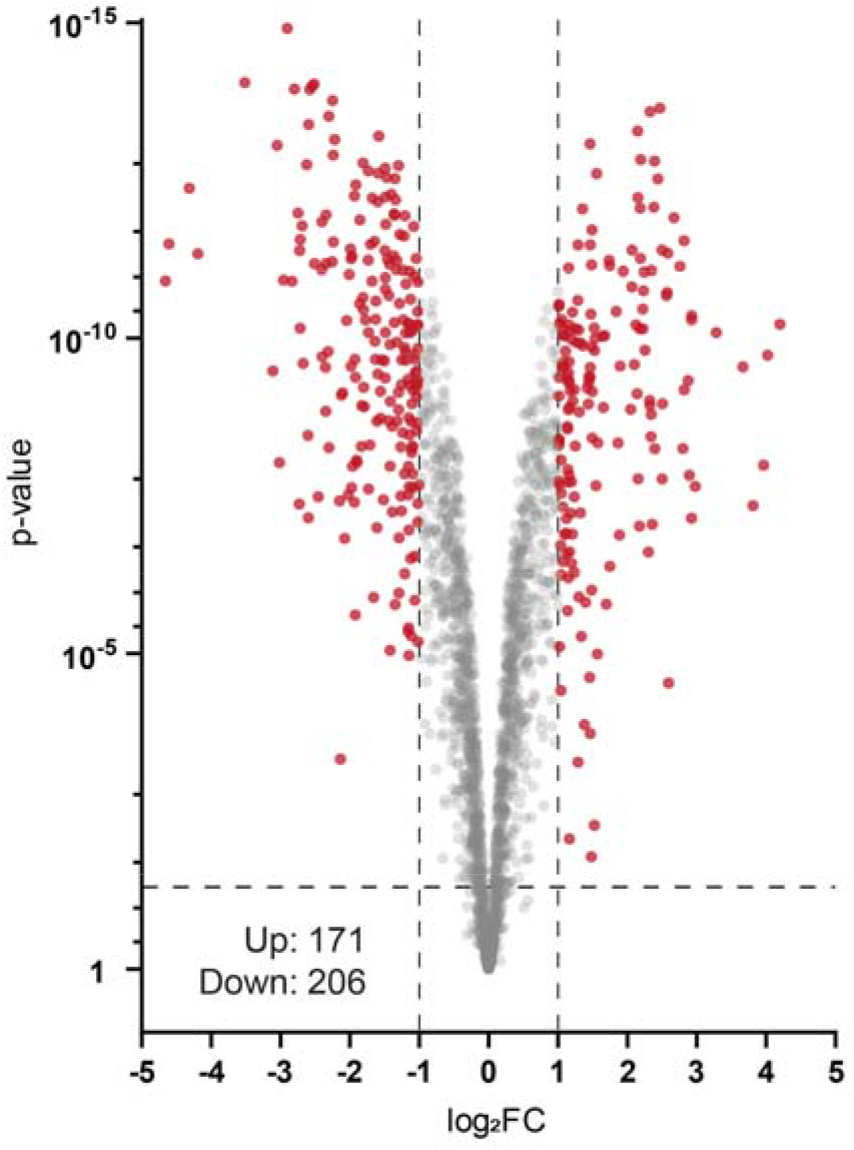
Volcano plot of differential protein levels in N-24 Δ*hfq* bacteria shown as a log_2_ change from wild-type bacteria. Proteins differentially expressed more than 1 log_2_ (i.e., a greater than 2- fold change) are coloured red. The number of differentially expressed proteins are indicated (log_2_FC > 1 and p-value <0.05).

